# On the problems of not accounting for measurement- and sampling uncertainty in ecological and environmental models

**DOI:** 10.1101/2020.03.16.993477

**Authors:** Christian Damgaard

## Abstract

In many applied cases of ecological and environmental modelling, there is a sizeable variation among the measured variables due to measurement- and sampling error. Such measurement- and sampling error among the independent variables may lead to regression dilution and biased prediction intervals in traditional empirical modelling. It is possible to avoid these shortcomings by subsampling and hierarchical modelling, but this is still not a common practice. Here, it is recommended to model the measurement- and sampling errors by integrating the stochastic modelling of the errors into hierarchical ecological and environmental models.

## Introduction

Predictions based on ecological and environmental models are often used in the counseling of the administrative and political system and thus play an important role for making political decisions on environmental issues (National Research Council 2007). Most directly, this is seen in fish stock models (Aeberhard et al. 2018), which have an immediate influence on the politically determined fish catch rates.

In order to strive for precision and realism, many ecological and environmental models have been developed to be functionally complex with many input variables and parameters (Evans 2012). This focus on realism has, in many cases, lead to downplaying the stochastic modelling of the data that are needed for fitting the models, and often the complex models are simply fitted to the observed mean values of the independent variables.

In traditional empirical modelling, it is assumed that the independent variables (or explaining variables) are known with no uncertainty (Fuller 1987). However, in many applied cases of ecological and environmental modelling there is a sizeable variation among the measured variables due to measurement- and sampling error (Yanai et al. 2018). For example, the soil type was determined with relatively large uncertainty in a study on wet heathlands where its effect on the vegetation was investigated (Damgaard 2019). Furthermore, some independent variables may covary. For example, in a climate experiment in a heathland ecosystem there was sizeable covariation between soil water and soil temperature, which had a significant effect on the inferred ecological conclusions of the experiment (Damgaard et al. 2018). Unfortunately, the measurement- and sampling errors are not always reported (Yanai et al. 2018).

Measurement- and sampling errors in the independent variables may seriously bias the estimated parameters and inferences in a statistical analysis (Fuller 1987, Chesher 1991). More recently, it has become possible to account for this variation and possible covariation among variables using hierarchical modelling methods (see Appendix A for a brief description of hierarchical models). However, these modelling methods are still not the preferred method of choice in the majority of the applied cases of ecological and environmental modelling.

The aim of this study is to use a simple linear model example for discussing the problem of falsely assuming that independent variables are known without any measurement- and sampling error with a special focus on the predictive precision of ecological and environmental modelling.

## Methods

One of the key assumptions in standard statistical modelling is that the independent variables are known with no uncertainty (Fuller 1987). For example, if the dependent variable, *y*, is modelled as a function of the dependent variable(s), *x*, then it is only the stochasticity of the dependent variable that is modelled,

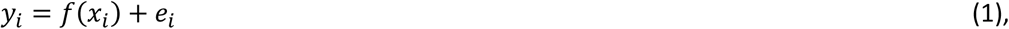

where the residual variation of the dependent variable is modelled by a suitable probability distribution. Often, the normal distribution is a suitable distributional choice, and the residual variation of observation *i* is then modelled by *e*_*i*_∼*N*(0, *σ*). However, in the following we will assume that there is a true, but unknown, value of the independent variable at observation *i, x*_*i*_, but that the observed value of the independent variable includes a measurement- and sampling error, *z*_*i*_, so that,

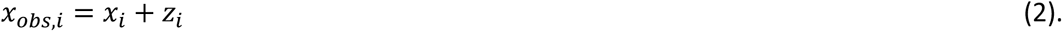

This measurement- and sampling error may have various case-dependent distributions, but here it will be assumed to be unbiased and normally distributed, i.e. *z*_*i*_∼*N*(0, *τ*), where *τ* is the standard deviation of the error. Furthermore, it is assumed that the value of the dependent variable at observation *i* is controlled by the true, but unknown, value of the independent variable, as described in eqn. 1.

A thousand datasets (*x*_*obs,i*_, *y*_*i*_) were simulated in a simple linear model, where *y*_*i*_ = *x*_*i*_ + *e*_*i*_, i.e. a linear model with an intercept of zero and a slope of one. Each dataset was fitted using linear models, and the mean estimates and prediction intervals (both mean and single-value prediction) of the resulting analyses are reported. The results of this simple measurement-error model are well-known in a general sense (Fuller 1987), but for researchers that are less familiar with the statistical literature, it is an advantage to consider more concrete examples and simulations.

For comparison, datasets generated under the same measurement-error model as above were fitted using Bayesian hierarchical models (Appendix A). That is, the joint posterior distribution of the true, but unknown, value of the independent variable at observation *i, x*_*i*_, was estimated together with the other model parameters using numerical MCMC methods (Metropolis algorithm). The prior distribution of the standard deviations was assumed to be inverse gamma distributed with parameters (0.1, 0.1), and the prior distribution of the location parameters was assumed to be improperly uniform distributed. A single chain of 100.000 iterations with a burn-in period of 50.000 was used to simulate the joint posterior distributions of the parameters. The marginal posterior distributions of the parameters as well as the posterior predictive distribution, *p*(*y*|*x* = 10), for both the mean and single-value predictions were summarized by their 95% credible intervals.

## Results

As expected (Fuller 1987), the estimated slope of the fitted linear model decreases, and the estimated intercept increases with measurement- and sampling error (Fig. 1, Table 1). This well-known phenomenon is known as “regression dilution” or “regression attenuation” in the statistical literature.

**Table 1.** Estimated mean properties of fitted linear models from 1000 realizations of the generated data model with an intercept of zero and a slope of one at different levels of measurement- and sampling error (*τ*). Sample size = 50, residual standard deviation (*σ*) = 1. The lower and upper 95% prediction intervals for both mean and single prediction intervals were made for *x* = 10, with an expected value of 10.

| $\tau$ | Intercept | Slope | Variance | Lower mean | Upper mean | Lower pred. | Upper pred. |
| --- | --- | --- | --- | --- | --- | --- | --- |
| 0 | -0.016±0.013 | 1.0031±0.0021 | 0.998±0.006 | 9.386±0.010 | 10.644±0.010 | 7.920±0.012 | 12.110±0.012 |
| 1 | 0.966±0.016 | 0.8315±0.0025 | 1.823±0.012 | 8.489±0.011 | 10.072±0.013 | 6.466±0.015 | 12.095±0.015 |
| 2 | 2.635±0.014 | 0.5433±0.0022 | 3.157±0.020 | 7.171±0.011 | 8.964±0.013 | 4.400±0.017 | 11.735±0.015 |
| 3 | 3.789±0.013 | 0.3410±0.0021 | 4.094±0.022 | 6.311±0.011 | 8.086±0.012 | 3.048±0.018 | 11.348±0.013 |

**Fig. 1.**
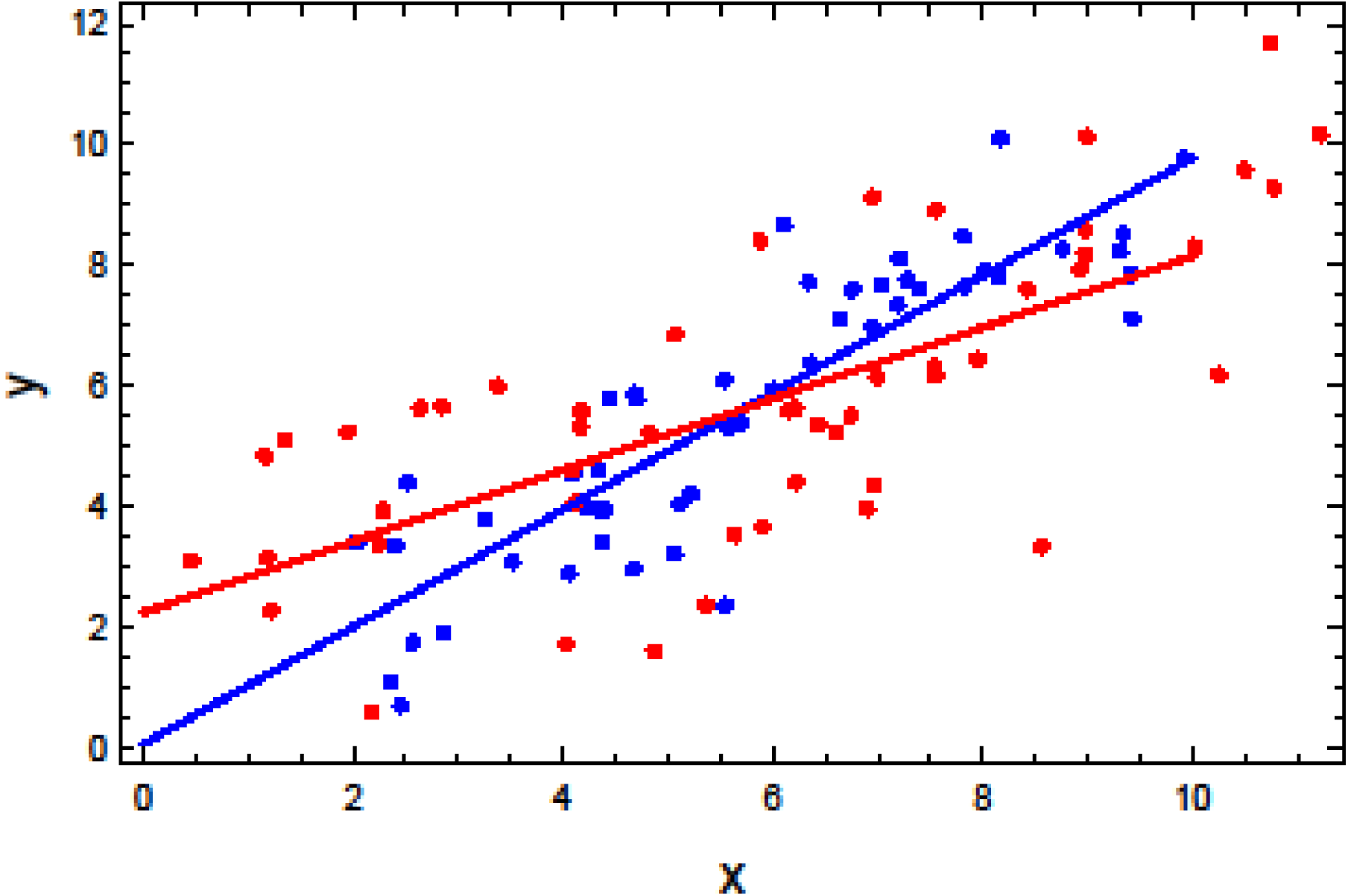
Two realizations of the generated data model with an intercept of zero and a slope of one at different levels of measurement- and sampling error, blue: *τ* = 0, red: *τ* = 2. The lines are the fitted lines. Sample size = 50, Residual standard deviation (*σ*) = 1.

Following directly from such regression dilution, it is clear that the predictive performance of the model is weakened at the borders of the domain of the independent variable. In the presented cases, the predictions at *x* = 10 (where the expected value of *y* = 10) may be heavily downward biased (Table 1). If the measurement- and sampling error was twice as large as the residual error (*τ* = 2), then the mean prediction interval was 7.17 - 8.96 and the single-value prediction interval was 4.40 - 11.74 (Table 1).

If the realizations of the data generating model were fitted using a Bayesian hierarchical model without subsampling, then the same regression dilution and downward biased prediction intervals were observed as when the data were fitted using the standard linear model (Table 2A). However, if each of the independent observations were subsampled ten times, i.e. each *x*_*i*_ is now determined by a sample of ten independent observations, then the Bayesian hierarchical model fitted the data adequately without any regression dilution or downward bias of the prediction intervals (Table 2B). This improved model fit was in this case mainly due to the subsampling and the consequently reduced sampling error and less due to the hierarchical modelling approach itself (Table 2C, where the hierarchical data structure was replaced by the mean of the subsamples). However, notice that the measurement- and sampling error was estimated correctly when the hierarchical data structure was modelled (Table 2B).

**Table 2.**
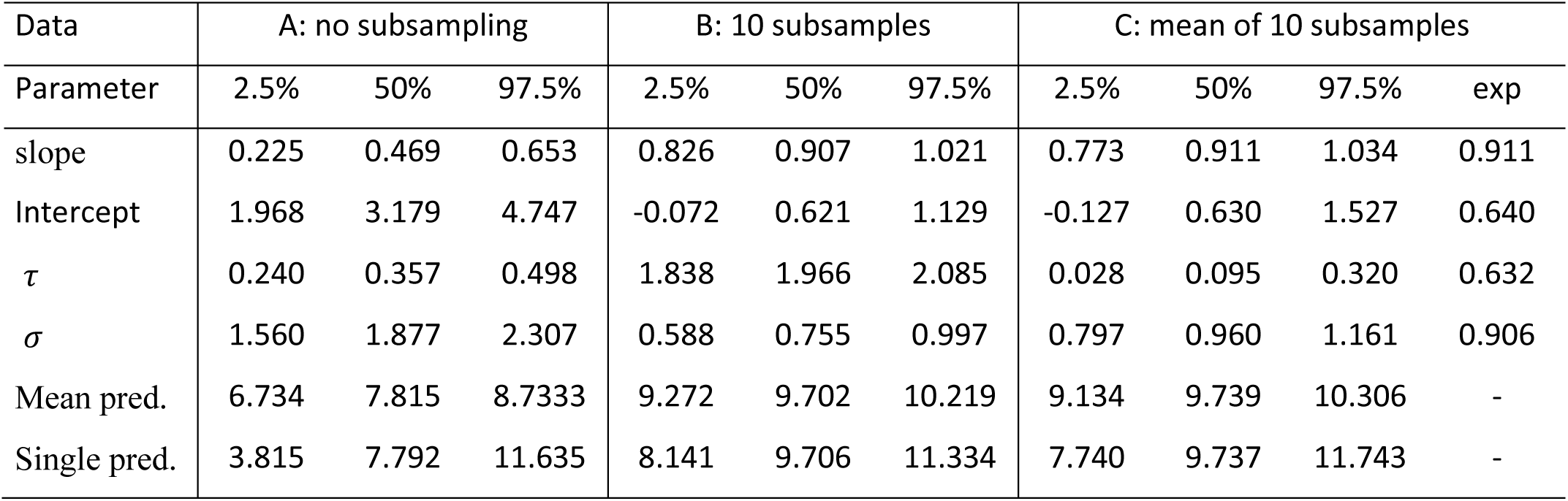
Summary of the marginal posterior distribution of a fitted linear model to one realization of the data generating model with an intercept of zero and a slope of one. Sample size = 50, measurement- and sampling error (*τ*) = 2, residual standard deviation (*σ*) = 1. Mean and single prediction intervals were made for *x* = 10, with an expected value of 10. A: no subsampling, B: each of the 50 independent values were subsampled 10 times, C: same as B, but the mean of the 10 subsamples were taken before fitting. For data realization B and C, the estimated values in a standard linear model are given in the column exp 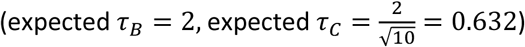.

| Data | A: no subsampling |  |  | B: 10 subsamples |  |  | C: mean of 10 subsamples |  |  |  |
| --- | --- | --- | --- | --- | --- | --- | --- | --- | --- | --- |
| Parameter | 2.5% | 50% | 97.5% | 2.5% | 50% | 97.5% | 2.5% | 50% | 97.5% | exp |
| slope | 0.225 | 0.469 | 0.653 | 0.826 | 0.907 | 1.021 | 0.773 | 0.911 | 1.034 | 0.911 |
| Intercept | 1.968 | 3.179 | 4.747 | -0.072 | 0.621 | 1.129 | -0.127 | 0.630 | 1.527 | 0.640 |
| $\tau$ | 0.240 | 0.357 | 0.498 | 1.838 | 1.966 | 2.085 | 0.028 | 0.095 | 0.320 | 0.632 |
| $\sigma$ | 1.560 | 1.877 | 2.307 | 0.588 | 0.755 | 0.997 | 0.797 | 0.960 | 1.161 | 0.906 |
| Mean pred. | 6.734 | 7.815 | 8.7333 | 9.272 | 9.702 | 10.219 | 9.134 | 9.739 | 10.306 | - |
| Single pred. | 3.815 | 7.792 | 11.635 | 8.141 | 9.706 | 11.334 | 7.740 | 9.737 | 11.743 | - |

## Discussion

The main conclusion of the simulations is that it is important to reduce any sizable measurement- and sampling error among the independent variables by independent subsampling of the variables. In principle, it is then possible to use traditional regression techniques to fit the mean values to the model. However, if the fitted modes have several independent variables with variable measurement- and sampling errors or if the sub-sampling is unbalanced, then some independent variables will be determined with larger precision than others, and it will be incorrect to use the means as the independent variables in empirical modelling. Instead, it is here recommended to *always* apply hierarchical modelling if there is any sizable measurement- and sampling error among the independent variables. In the cases, where it is unknown whether there is a sizable measurement- and sampling error among the independent variables then it is recommended to investigate this in a sensitivity analysis by simulating *df*(*x*)/*d z*.

Furthermore, in the performed simulations the distribution of the measurement- and sampling error was assumed to be independent, unbiased and symmetric. However, such regularity assumptions may not always be adequate in ecological and environmental models, and then it becomes even more important to model the distributional properties of the measurement- and sampling error in hierarchical models. For example, plant cover data of dominating species may be either L or U-shaped due to the spatial aggregation of plant species, and it has become common to model the measurement- and sampling error of plant cover of single species by the beta distribution (Damgaard 2009, Damgaard and Irvine 2019).

Additionally, the measured variables may covary, and it will be incorrect not to consider this covariation during model development and fitting. For example, there is an increasing awareness that it is important to model the joint species distribution in ecosystem models instead of treating each species as being independent (Ovaskainen et al. 2017), and e.g. plant cover data, which often are found to be strongly negatively correlated, have been suggested to be modelled by the Dirichlet distribution (Damgaard 2015, 2018). The omission of important variable covariation in model development and fitting may lead to strongly biased parameter estimates and prediction intervals in specific cases (Damgaard et al. 2018), and it seems reckless *a priori* to neglect the effect of such covariation due to a preferred use of non-hierarchical modelling techniques.

In order to strive for precision and realism, many ecological and environmental models have been developed to be complex with many input variables and parameters (Evans 2012). However, it is important that this model complexity does not hinder a proper stochastic modelling of the data that are used to fit the model. As demonstrated here, this may lead to regression dilution and biased prediction intervals. Fortunately, hierarchical modelling has become practically easier to perform and there are now appropriate software solutions that eliminate the need to make this compromise.

### Appendix: Hierarchical modelling

Hierarchical models (also called latent variable- or state-space models) have received considerable attention as a flexible and powerful tool to fit models to ecological data (Clark 2007, Ovaskainen et al. 2017, Aeberhard et al. 2018, Damgaard 2019). A hierarchical model consists of measurement equations that link the sampled data to the underlying features of the system that need to be modelled (Fig. A1). These features, which are called latent variables, may simply represent the unknown mean of a sampled variable and, in this case, the measurement equation models the sampling variance of the observed data. The studied processes are modelled by process equations, where the causal relationships between the latent variables are modelled (Fig. A1). Typically, the causal relationships are only partly known, and the remaining uncertainty is modelled as stochastic structural uncertainty.

In a standard simple regression model of an independent and a dependent variable, the independent variable is assumed known without error and only one type of variance is estimated, the residual variance. However, in the simple hierarchical model shown in Fig. A1 the variance is partitioned into two sampling variances and a structural variance, where the latter is particularly relevant when making predictions.

In a Bayesian framework, all the arrows may be modelled as conditional probabilities and assuming conditional independence, i.e. the child node depends only on the parent node(s), then the likelihood function of the whole system may be specified by multiplying the likelihood functions of all the conditional probabilities. The resulting combined likelihood may be complex, and the joint posterior distribution of the model parameters and the latent variables are typically calculated using numerical Markov chain Monte Carlo (MCMC) methods.

If the data are hierarchical, e.g. if there is a spatial and/or temporal structure in the data, then this random-effect structure may conveniently be modelled by introducing new latent variables that mimic the hierarchical structure of the data, e.g. by introducing a new latent variable for each site and/or year. Compared to mixed models, these latent variables may be assumed to arise from a common distribution by introducing hyper-priors (Clark 2007, Gelman and Hill 2007).

**Fig. A1.**
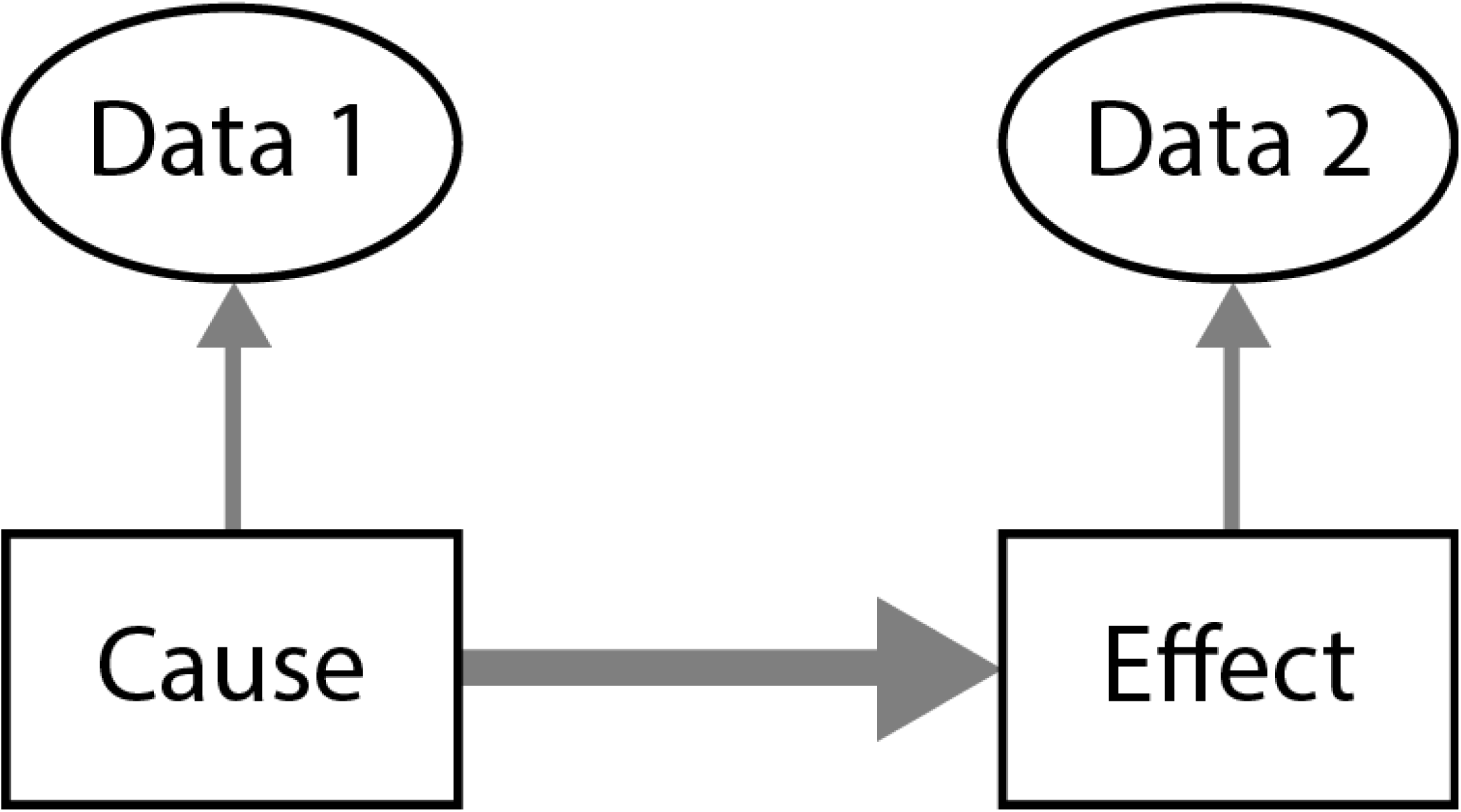
Outline of a simple hierarchical model. Latent variables are shown as rectangles, and sampled data are shown as ovals. The thin arrows denote measurement equations, and the thick arrow denotes the process equation.

